# WormCat 2.0 defines characteristics and conservation of poorly annotated genes in *Caenorhabditis elegans*

**DOI:** 10.1101/2021.11.11.467968

**Authors:** Daniel P. Higgins, Caroline M. Weisman, Dominique S. Lui, Frank A. D’Agostino, Amy K. Walker

**Affiliations:** Program in Molecular Medicine, UMASS Chan Medical School, Worcester MA; Lewis-Digler Institute for Quantitative Genomics, Princeton University, Princeton, NJ; Department of Applied Mathematics, Harvard University, Cambridge MA

## Abstract

Genome-wide measurement of mRNA or protein levels provides broad data sets for biological discovery. However, subsequent computational methods are essential for uncovering the functional implications of the data as well as intuitively visualizing the findings. Current computational tools are biased toward well-described pathways, limiting their utility for novel discovery. Recently, we developed an annotation and category enrichment tool for *Caenorhabditis elegans* genomic data, WormCat, that provides an intuitive visualization output. Unlike GO, which excludes genes with no annotation information, WormCat 2.0 retains these genes as a special UNASSIGNED category. Here, we show that the UNASSIGNED gene category enrichment exhibits tissue-specific expression patterns and include genes with biological functions. Poorly annotated genes have previously been considered to lack homologs in closely related species. Instead, we find that around 3% of the UNASSIGNED genes have poorly characterized human orthologs. These human orthologs are themselves have little annotation information. A recently developed method that incorporates lineage relationships (abSENSE) indicates that failure of BLAST to detect homology explains the apparent lineage specificity for many UNASSIGNED genes, suggesting that a larger subset could be related to human genes. WormCat provides an annotation strategy that allows association of UNASSIGNED genes with specific phenotypes and known pathways. Our analysis indicates that the UNASSIGNED gene category contains candidates that merit further functional study which could yield insight into understudied areas of biology.

## Introduction

Unbiased assays such as genetic screens, transcriptomics, and proteomics are powerful tools for identifying genes that are critical players in developmental processes, regulated in response to stress, or altered in disease processes. Recent technological improvements have vastly improved data quality and decreased the cost of transcriptomic analysis. Deep sequencing of mRNAs (RNA-seq) has, therefore, become a widely used assay to compare gene expression across cell types in differing physiological conditions or mutant backgrounds (Li et al., 2014). Proteomics approaches are also increasingly common and sophisticated, and allow quantitative analysis of peptides present in specific subcellular compartments or carrying post-translational modifications (Fonslow et al., 2014). The genetic tools available in *C. elegans* complement these assays, as the results of -omics experiments can readily be subjected to functional validation and follow up analyses. However, genome-wide studies have limitations as discovery tools. Unlike classical genetic approaches that allow the study of genes before the functions are known, -omics experiments depend on pathway analysis that direct focus toward well-studied genes and pathways.

While -omics technologies can generate large amounts of high-quality data, several challenges complicate data interpretation, gene categorization, and data visualization. GO (Gene Ontogeny) classifications are a broadly used platform that annotates data from -omics analysis according to biological or molecular function and cellular location. This analysis returns categories that are statistically enriched in the input relative to the entire genome (Angeles-Albores et al., 2018a; Eden et al., 2009; Mi et al., 2019). Other gene annotation databases, such as KEGG, provide pathway or enzymatic functional data (Thomas, 2016). While these resources contain valuable data, the highest-quality information is found for the most highly studied genes (Wood et al., 2019); thus, they are of maximal utility in pathways that are already well-studied. In addition, genes with unclear functions are often excluded from enrichment analysis (Ding et al., 2018). Genome-wide analysis also generate large and complex datasets, making intuitive data visualization an additional challenge. While bar charts that graph p-values are useful for visualizing single data sets, other styles such as bubble charts may be more useful for comparing data across multiple conditions.

While *C. elegans* is amenable to genome-wide assays and multiple tools exist for pathway analysis of the results, most depend on GO annotation (Angeles-Albores et al., 2018b; Eden et al., 2009; Mi et al., 2019). However, we previously noted that commonly used GO servers excluded around 30% of *C. elegans* genes and that many GO categories poorly describe gene function (Ding et al., 2018). To circumvent these issues, we developed WormCat, a web-based program based on a near-complete annotation of the *C. elegans* genome (Holdorf et al., 2019). WormCat provides annotation for each input gene, determines category enrichment within the gene set, and provides scaled bubble charts for visualization. In the original version of WormCat, we included a category for poorly annotated genes (UNKNOWN) to avoid biasing the annotation lists toward well-studied genes. Using the whole genome annotation list, we found that WormCat identified biologically significant categories from *C. elegans* exposed to RNAi or pharmacological treatments as well as from tissue-specific RNA-seq studies (Holdorf et al., 2019). In addition, annotation lists specific for the commonly used RNAi libraries revealed enriched pathways in data from an RNAi screen. WormCat has been rapidly adopted by the *C. elegans* community (Albarqi and Ryder, 2021; Lee et al., 2021; Naim et al., 2021; Tecle et al., 2021; Zhang et al., 2021) and adapted into a metabolism-focused tool (Walker et al., 2021) as well as an integrated gene expression analysis program (Cheng et al., 2021).

Category enrichment tools provide a valuable method for identifying patterns in complex data sets. However, as well-studied genes and pathways are often the best annotated, genes that are poorly annotated are less likely to be chosen for further analysis. Thus, pathway analyses tools bias insights gained from -omics analysis toward what is already known, leaving a pool of potentially important genes consistently unanalyzed by studies. Here we focus on developing the UNASSIGNED category in WormCat to allow examination of poorly annotated genes to stimulate future functional studies. We find that representation of UNASSIGNED genes differs across tissues in published RNA-seq datasets. In addition, this category is poorly represented in multiple whole-animal proteomics datasets. By identifying enrichment and expression characteristics of genes with poorly defined functions, WormCat 2.0 will stimulate the inclusion of previously under-analyzed genes in functional studies. As 3% of poorly annotated genes have human orthologs, modeling analysis of unassigned genes in *C. elegans* provides a roadmap that can be used to extend functional analysis of these genes. In addition, our analysis provides impetus to revisit poorly annotated gene across model organisms.

## Results

### Evaluation of UNKNOWN/UNASSIGNED genes in WormCat 2.0

WormCat is a program for pathway analysis of *C. elegans* RNA seq, ChIP seq, or genetic screen data that utilizes a near-complete annotation list of *C. elegans* genes (Holdorf et al., 2019). Input genes (Regulated Gene Sets, RGS) entered as a single list or in a batch file are mapped to the specific annotation lists. Fisher’s Exact Test is used to determine the statistical significance of enrichment by building a contingency table that compares the number of genes in each category in the RGS to the entire annotation list along with a false discovery rate correction (See Figure 1A). Because RNA-seq or ChIP-seq data contain information on non-coding RNA expression, the whole genome annotation includes lincRNAs, miRNAs, snoRNAs, tRNAs, and pseudogenes. To enable category enrichment analysis of RNAi library screens, WormCat includes RNAi library-specific annotation lists to correct for the sub-genomic size of the RNAi libraries (Holdorf et al., 2019). In addition, WormCat 2.0 provides an appropriate background ORF-only (open read frame) option for proteomics data sets (Figure 1A). Category enrichment results from WormCat are provided as a scaled bubble chart (.sgv), a sunburst diagram, and .csv files. To facilitate the comparison of multiple data sets, we have added a batch processor also which produces a combined output file.

**Figure 1:**
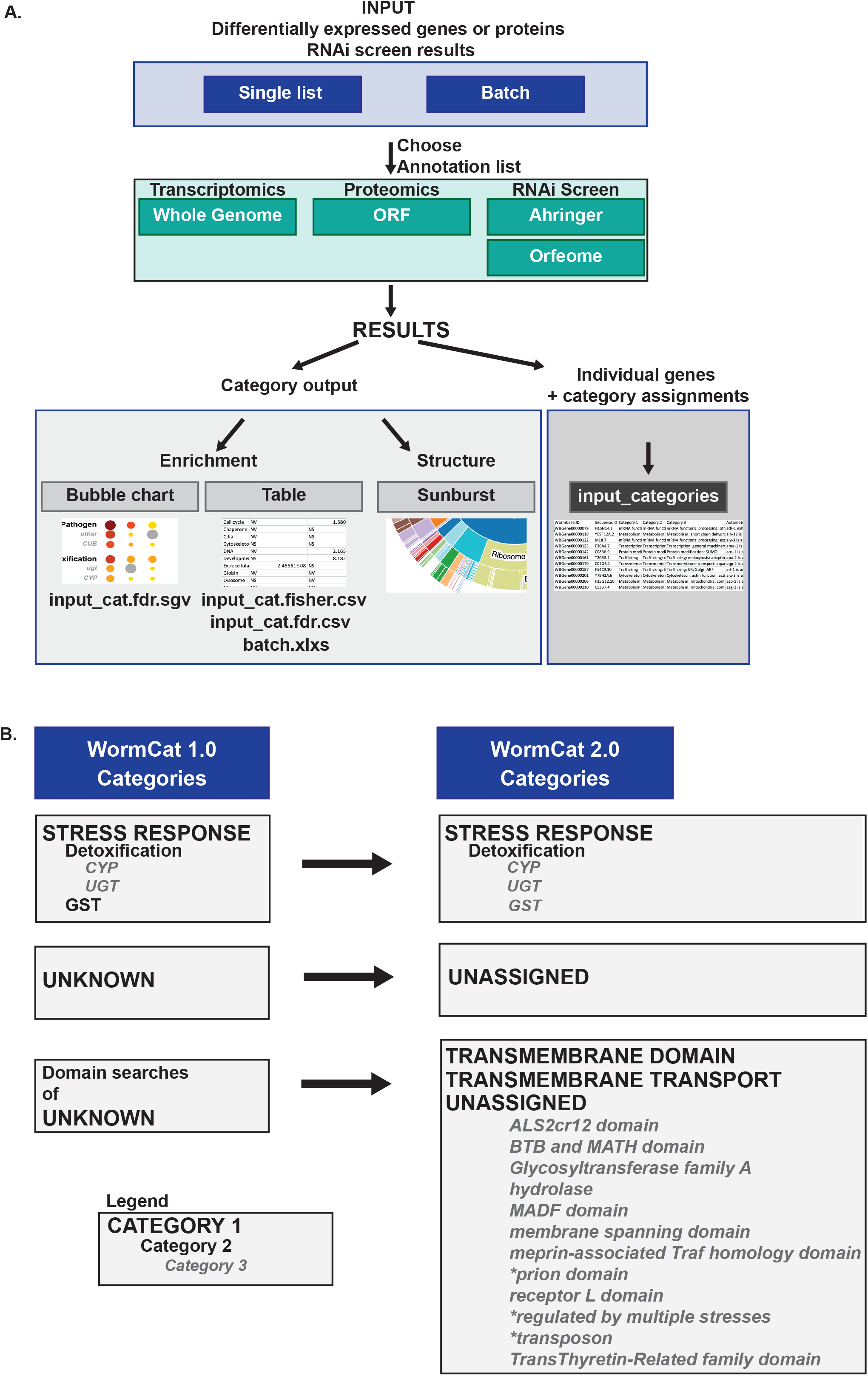
WormCat 2.0 supports category enrichment of multiple -omics datasets and allows identification of less well-characterized genes. **A**. Schematic diagram of WormCat 2.0 workflow. **B**. Changes in the annotation list structure from WormCat 1.0 to WormCat 2.0. See **Table S1** for annotation of each *C. elegans* gene, **Table S2** for annotation definitions and **Table S3** for the list restricted to protein-coding genes. Abbreviations: TM, transmembrane; CYP, cytochrome p450; CUB, complement C1r/C1s, Uegf, Bmp1; ugt, uridine diphosphate glucuronosyl transferase; GST, Glutathione S-transferase.

The WormCat whole genome annotation list contains 31,390 genes divided into nested categories, with 35 Category 1 (Cat1) groups divided into 242 Cat3 and 361 Cat3 designations. Categories are based on physiological functions. If the biological role was unclear or pleiotropic, genes were placed in molecular categories or annotated based on localization (Holdorf, et al. 2019). Division of Cat1 into Cat2 and Cat3 groupings also depended on biological or molecular function, then localization. For example, in the “STRESS RESPONSE” category, we noted that Glutathione-S-Transferases (GSTs) were placed at a Cat2 level. In contrast, other enzymes involved in response to xenobiotics were in STRESS RESPONSE: Detoxification. We therefore reassigned the GSTs (Figure 1B, Supplementary Tables 1, 2).

Gene annotation and gene set enrichment within genome-scale experiments are often used to select pathways or genes for further analysis. However, genes that are poorly annotated or appear to be lineage-specific may not be prioritized. While human orthology may be important for certain *C. elegans* studies, some of these genes may have homology not recognized by sequence comparison algorithms (Weisman et al., 2020). Even if genes are lineage-specific, they may have critical biological functions. Therefore, a comprehensive analysis requires annotation criteria from publicly available sources that will allow subsequent characterization of these genes. In the initial version of WormCat, we placed genes that did not meet annotation criteria for other categories in the “UNKNOWN” category so that statistical calculations included all genes, regardless of functional annotation. This category is now labelled “UNASSIGNED” to better reflect the common characteristic of this gene group (Supplemental Tables 1, 2). Within Cat2 or Cat3, less well-characterized genes were also changed from “Cat2:other” to “Cat2: unassigned”. In the initial WormCat annotation list, 8160 genes were classified as UNKNOWN, representing 26% of the *C. elegans* genome. Many of these genes had WormBase annotations with cellular locations or protein domains of unclear function (Harris et al., 2019). In our initial WormCat annotation list, we noted prion domains or induction of expression in response to multiple stresses at the Cat3 level. To expand these annotations, we used the KEGG database (Kanehisa and Goto, 2000), NCBI Conserved Domain Database (Lu et al., 2019), TOPCONs membrane domain (Bernsel et al., 2009), and UNIPROT cellular localization predictions (Consortium et al., 2020)to examine each gene in the UNASSIGNED category (Supplemental Table 4). As a result, 81 genes were reassigned into Cat1 groupings such as “METABOLISM,” “STRESS RESPONSE,” or mRNA FUNCTIONS” based on close inspection of homology-based UNIPROT annotation, or subcellular localization.

WormCat 1.0 placed membrane-spanning proteins with well-described functions into biological or molecular-based categories. Genes encoding transmembrane regions (TMs) identified by the NCBI conserved domain database were placed in the “TRANSMEMBRANE DOMAIN” group (Holdorf et al., 2019). However, this excluded many genes with WormBase descriptions including the phrases “localizes to plasma membrane” or “ER protein” (Harris et al., 2019). To more rigorously identify transmembrane proteins, we ran all the genes in the “UNASSIGNED” and “TRANSMEMBRANE DOMAIN” categories in the TOPCONS suite, which includes six algorithms for transmembrane domain or signal sequence identification (Bernsel et al., 2009). We used the following criteria to assign categories (Figure 1B, Supplementary Table 4). First, 41 genes with TM domains in all TOPCONs algorithms and domains common to transporters were reassigned to “TRANSMEMBRANE TRANSPORT.” Next, 1466 genes showing predicted transmembrane regions with all six programs in the suite, but lacking transporter-associated domains were placed in “TRANSMEMBRANE DOMAIN” (Figure 1B, Supplemental Table 4). Next, those with transmembrane regions predicted in one or two of the TOPCONS programs were placed in the UNASSIGNED: Unassigned: membrane-spanning domain. Finally, proteins with a predicted signal sequence but no TM domain were assigned to Extracellular material: secreted protein. Analysis of the TRANSMEMBRANE DOMAIN proteins with TOPCON showed that 32 did not have 6/6 predictions from TOPCONS and were therefore moved to UNASSIGNED: Unassigned: membrane-spanning domain to denote lower confidence scores (Figure 1B; Supplemental Table 4).

Some domain names, such as “BTB-POZ” or “Receptor L domain” identified by GO or the Conserved Domain Database, do not have precise functional characterizations, and others such as “hydrolase” may reflect a general enzymatic function, but the appropriate category is unclear. Therefore, we added nine new Cat3 level annotations for multiple common but functionally unclear domains in the UNASSIGNED category. Based on these re-annotations, 2251 genes in UNASSIGNED were assigned to other categories or given additional category information at the Cat3 level. These annotation reassignments both improve the accuracy of WormCat and increase the curated information that can be applied to the UNASSIGNED genes.

### Levels of lineage specificity in the UNASSIGNED gene category

One goal of providing curated information on UNASSIGNED category genes is to allow WormCat users to identify those with potential biological functions based on expression in RNA-seq, proteomic, or RNAi screen data sets. We next analyzed how many UNASSIGNED genes are annotated in GO, have human orthologs, or appear in lineage-specific gene families. Many unannotated genes are excluded from commonly used web-based GO servers (Ding et al., 2018; Holdorf et al., 2019). In order to determine if the WormCat UNASSIGNED category overlaps with unannotated *C. elegans* genes in GO, we searched GO terms for each protein-coding WormBase ID in WormCat using the Parasite Biomart (Howe et al., 2017). We found that while 26% of protein-coding genes in WormCat lack associated GO terms, 72% of genes in the UNASSIGNED category are not annotated in GO. This makes the UNASSIGNED category the largest category of *C. elegans* genes that are not represented in GO (Figure 2A, B, Supplemental Table 4). Other WormCat categories with significant numbers of non-GO annotated genes include STRESS RESPONSE (11%), PROTEOLYSIS PROTEOSOME (20%), and NUCLEIC ACID (19%) (Figure S1A, Supplemental Table 4). Within the UNASSIGNED category, we annotated genes that were regulated in response to multiple stresses (MSR), contained a predicted transmembrane domain, or domains of unclear function (Holdorf et al., 2019), see also Fig 1B). While some of these categories are described in GO (membrane-spanning, TTR, hydrolase, and BTB/MATH), our annotation strategy separates higher or lower confidence transmembrane proteins and allows identification of the MSR genes along with additional shared domain proteins (Figure 2C, Supplemental Table 4).

**Figure 2:**
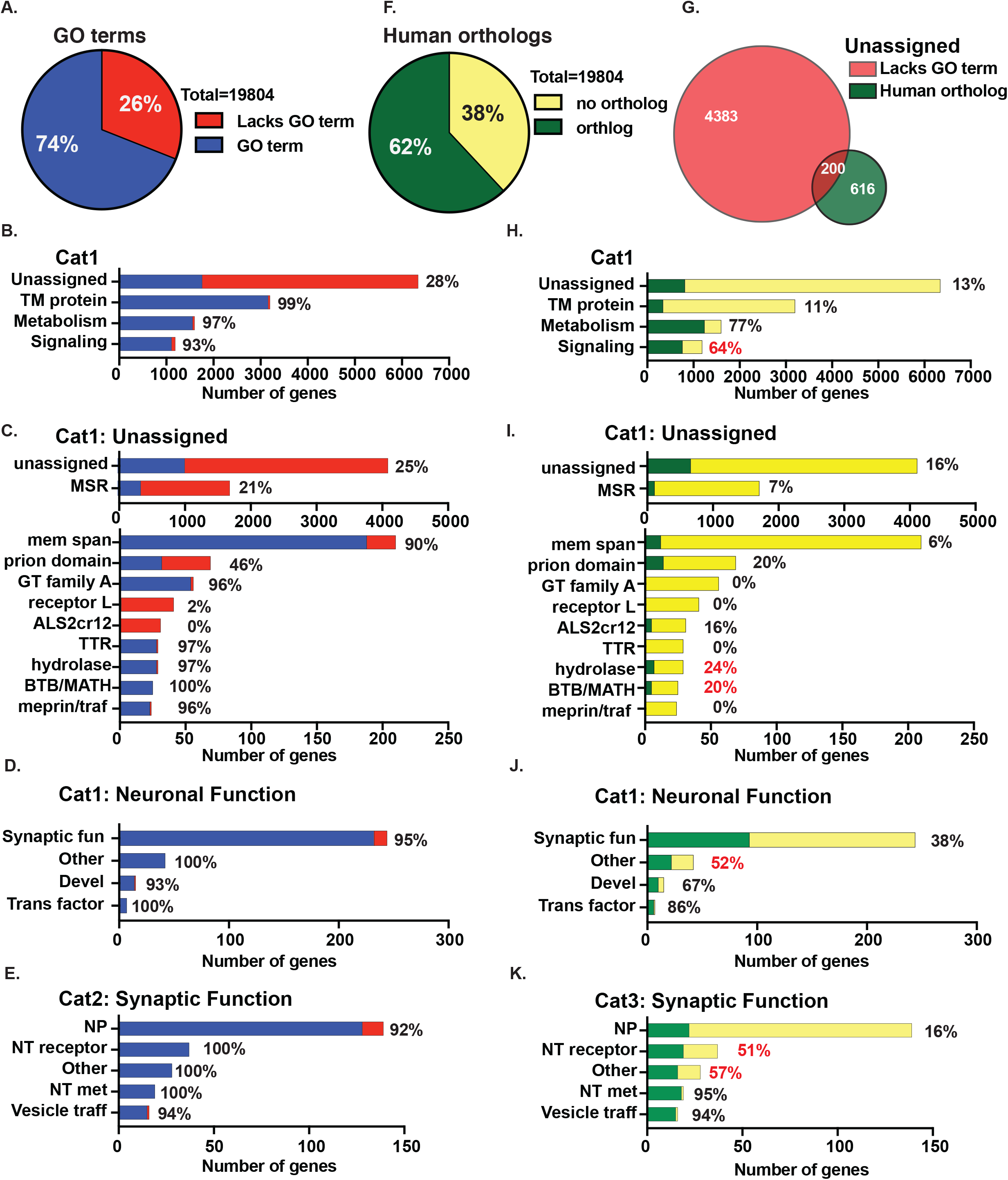
Unassigned genes in *C. elegans* include a subset with human orthologs. **A**. Pie chart of *C. elegans* protein-coding genes that are assigned GO terms in the ParaSite Biomart. **B**. Breakdown of WormCat Cat1 level categories with numbers of genes annotated by GO. Cat2 GO breakdown of Unassigned (**C**) and Neuronal function (**D**) with Cat3 level categories of Synaptic Function. Additional categories are in **Supplemental Figure 1 A-D. F**. Pie chart of *C. elegans* protein-coding genes designated as having human orthologs in the ParaSite Biomart. **G**. Venn diagram showing the overlap between Unassigned genes lacking GO terms and those with human orthologs. **H**. Breakdown of WormCat Cat1 level categories with numbers of genes with human orthologs. Cat2 human ortholog breakdown of Unassigned (**I**) and Neuronal function (**J**) with Cat3 level categories of Synaptic Function (K). **Supplemental Figure 1 E-H** Red percentages denote categories with substantially more human orthologs in a genome-wide BLASTP comparison of each *C. elegans* gene with the human genome (see **Table S4**). Abbreviations: TM, transmembrane; msr, multiple stress-regulated; mem span, membrane-spanning; GT family A, glucosyltransferase family A; TTR, TransThyretin-Related family domain; BTB/MATH, BR-C, ttk, and bab/meprin and TRAF homology; Synaptic fun, Synaptic function; Devel, Development; Trans, Transcription; NP, Neuropeptide, NT receptor, Neurotransmitter receptor; NT met, Neurotransmitter metabolism; Vesicle traff, Vesicle traffic.

Genes described as having human orthologs may be more extensively studied and better annotated (Wood et al., 2019). In order to determine if WormCat categories of genes lacking GO annotations were also predicted to lack human orthologs, we used the Parasite Biomart to obtain human ortholog predictions for each WormCat protein-coding gene. This website identified human orthologs for 62% of the *C. elegans* genes (Figure 2F, Supplemental Table 4), and is referrenced to the current WormBase release in comparison to Ortholist (Kim et al., 2018). We noted that WormCat categories that were poorly annotated by GO also had low percentages of human orthologs (Figure 2B, C, H, I, Supplemental Table 4). However, human orthologs were present in these categories, for example, the UNASSIGNED category contains 816 genes with human orthologs, 200 of which also lack a GO term (Figure 2G). This analysis indicates that genes within the UNASSIGNED category have been understudied even though some have predicted orthology to human genes.

Other categories such as TM protein and STRESS RESPONSE were relatively well-annotated by GO but had fewer predicted human orthologs in the Parasite database (Figure 2B, H; Supplemental Figure 2A-C; E-G; Supplemental Table 4). As an alternative method of determining potential orthology, we performed BLASTP on each gene in WormCat. We found that around 2800 genes that lacked human orthology annotation by Parasite Biomart had statistically significant e-values and had bit scores above 40 (Supplemental Table 4), suggesting that some of these genes could have human orthologs. However, the genes that were absent from the Biomart orthology database were largely from expanded gene families and were only 7% of the UNASSIGNED category (Figure 2 I-K, Supplemental Table 4).

Many core-biological function genes are well conserved across phyla (Chervitz et al., 1998) and are extensively characterized. Sequence conservation between proteins is often determined by pairwise alignment determined by BLAST (Altschul et al., 1990) or with algorithms such as HMMR (www.hmmer.org). Genes that lack detectable homology by these methods are referred to as “lineage-specific,” often with the implication that they have specalized function (Cai et al., 2006). However, some of these genes may have structural conservation despite lacking sequence homology detectable by BLAST (Pearson, 2013). Other genes may indeed encode proteins with lineage-specific functions or belong to classes of genes undergoing rapid evolution (Cai et al., 2006). For example, pathogen response genes may evolve rapidly to balance selection pressure (Sironi et al., 2015). In order to compare the number of lineage-specific genes in the UNASSIGNED category with other categories, we determined the number of species-specific and genus-specific genes (Zhou et al., 2015) defined by the absense of detected homologs in more distant species in each category (Figure S2A-H, Supplemental Table 4). We found that many categories had percentages of non-lineage specific genes that were close to numbers of human orthologs (Figure 2H-K, Figure S2A-H, Supplemental Table 4), as might be expected. About half of the UNASSIGNED genes were found by Zhou et al. to be lineage-specific, with similar proportions in subcategories of completely undescribed genes or within the MSR subcategory (Figure S2C). Domain-defined subcategories had fewer lineage-specific genes. Lineage-specific genes were largely contained within the UNASSIGNED genes lacking human orthologs, defined by the Parasite Biomart (Figure S2D). However, our BLASTP results suggest some of these genes (bit score >40 and an e value less than 0.01) contain sequence similarity to human genes (Figure S2J, Supplemental Table 4).

It is also possible that proteins have evolutionarily conserved functions, but the amino acid similarity is not apparent by BLAST or other algorithms (Pearson, 2013). A recent study has developed a phylogenetic method (abSENSE) to assess whether a gene could be lineage-specific for this reason. This method uses distance matrixes to predict the likelihood that orthologs in an outgroups could be undetectable by BLAST (Weisman et al., 2020). Using yeast and *Drosophila*, this study found that many genes considered to be lineage-specific arise from a failure to detect homology using present methods and also identified gene homologies that require an explanation beyond homology detection failure. This latter class were highlighted as particularly interesting candidates for functional studies. We adapted this method to estimate if any of the genes in the UNASSIGNED category could have undetected orthologs outside *Caenorhabditis* compared to the lineage-specific genes defined by Zhou et al. abSENSE makes predictions for whether a gene may have a homolog in a target species based on the evolutionary distances between that target species and the focal species (Weisman et al., 2020). We chose to include *C. elegans* and target species comprising two sister species, *C. remenai*, and *C. briggsae*, along with Clade III nematodes, *Necator americanus, Loa loa*, and *Brugia maylai*, and so calculated evolutionary distances between these species (Figure 3A, B). We also included a more distantly related invertebrate, the sea urchin *Strongylocentrotus purpuratus*. Most UNASSIGNED genes had orthologs in some of these target species, suggesting that they are not specific to *Caenorhabditis* (example in Figure 3C, Supplemental Table 4). Other genes were specific to *Caenorhabditis*; among these, we identified cases in which any potential homologs outside of the genus are likely to be undetectable (Figure 3D). For these genes, homologs may indeed be present in outgroup species: in this scenario, they appear lineage-specific merely because these homologs have diverged too far to be detected by standard homology search. Finally, we also identified cases in which potential homologs outside the genus, specifically in *S. purpuratus*, would likely have been detected if present, implying that they may be truly specific to the *Caenorhabditis* genus. We used WormCat to classify these potentially novel genes and found that the majority were in the UNASSIGNED, TRANSMEMBRANE: 7TM and PROTEOLYSIS: E3:Fbox family (Figure 3E, Supplemental Table 4), which are examples of rapidly expanding gene families in *C. elegans* (Robertson and Thomas, 2006).

**Figure 3:**
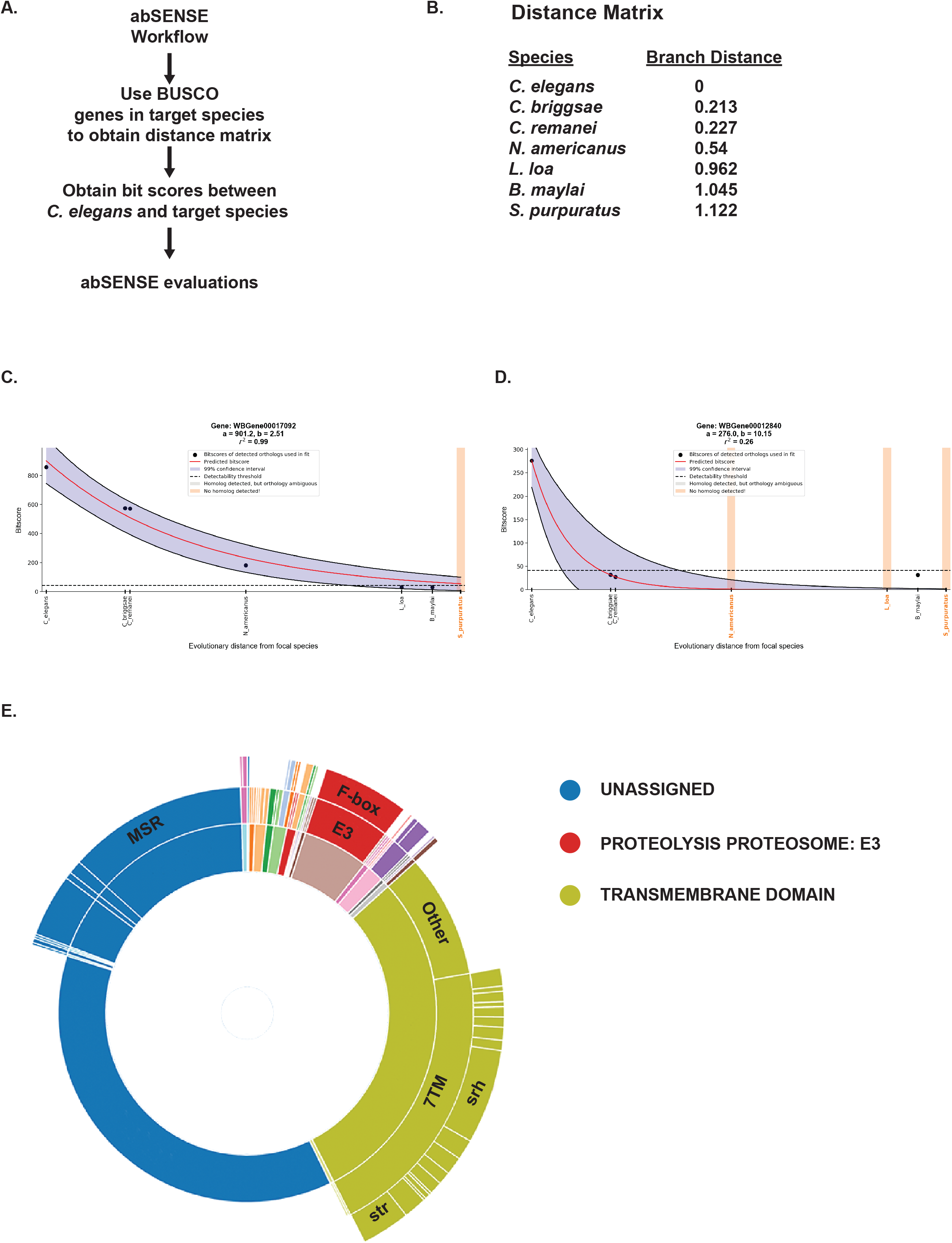
abSENSE analysis find many lineage-specificity of UNASSIGNED could be overestimated. **A:** Schematic showing abSENSE workflow. **B**. Distance matrix of species used in analysis. Example Venn diagrams (**C, D**). **E**. WormCat sunburst showing categories of *C. elegans* genes unlikely to have homologs in *S. purpuratus*.

### UNASSIGNED: regulated by multiple stress genes show *pmk-1* and *atf-7*-dependence

Annotation changes in WormCat 2.0 altered multiple categories, including the STRESS RESPONSE, TRANSMEMBRANE PROTEIN, TRANSMEMBRANE TRANSPORT, and UNASSIGNED categories. In order to test how WormCat2.0 performs, we determined gene enrichment on published RNA-seq data that examined which *Pseudomonas aeruginosa* (PA14) upregulated genes were also dependent on the MAP Kinase *pmk-1*/MPK13 and the bZIP transcription factor *atf-7* (Fletcher et al., 2019) (Figure 4A). Both WormCat 2.0 and 1.0 show highly significant enrichment in STRESS RESPONSE: Pathogen and Detoxification (Figure 4B, C; Supplemental Table 5), consistent with the authors’ GO enrichment findings as well as previous results (Ding et al., 2018; Fletcher et al., 2019; Troemel et al., 2006). The STRESS RESPONSE: Detoxification category has a slightly higher enrichment in WormCat 2.0 since GSTs are included at the Cat3 level (Figure 4D, E). Reannotation of the UNASSIGNED genes also added genes to the TM TRANSPORT category. However, we do not find significant changes in enrichment in TM TRANSPORT in this gene set from WormCat 1.0 to 2.0 (Figure 4A, B; Supplemental Table 5). The “UNASSIGNED: regulated by multiple stresses” category is enriched in *C. elegans* exposed to PA14 in this study and a previous version using WormCat (Ding et al., 2018; Fletcher et al., 2019) (Figure 4B, C; Supplemental Table 5). The UNASSIGNED: regulated by multiple stresses genes (MSR genes) lack characteristics that allow assignment into physiological or molecular categories but had changes in expression due to at least two common stress agents such as paraquat, methylmercury, or tunicamycin (see Holdorf et al. 2020, for a complete list).

**Figure 4:**
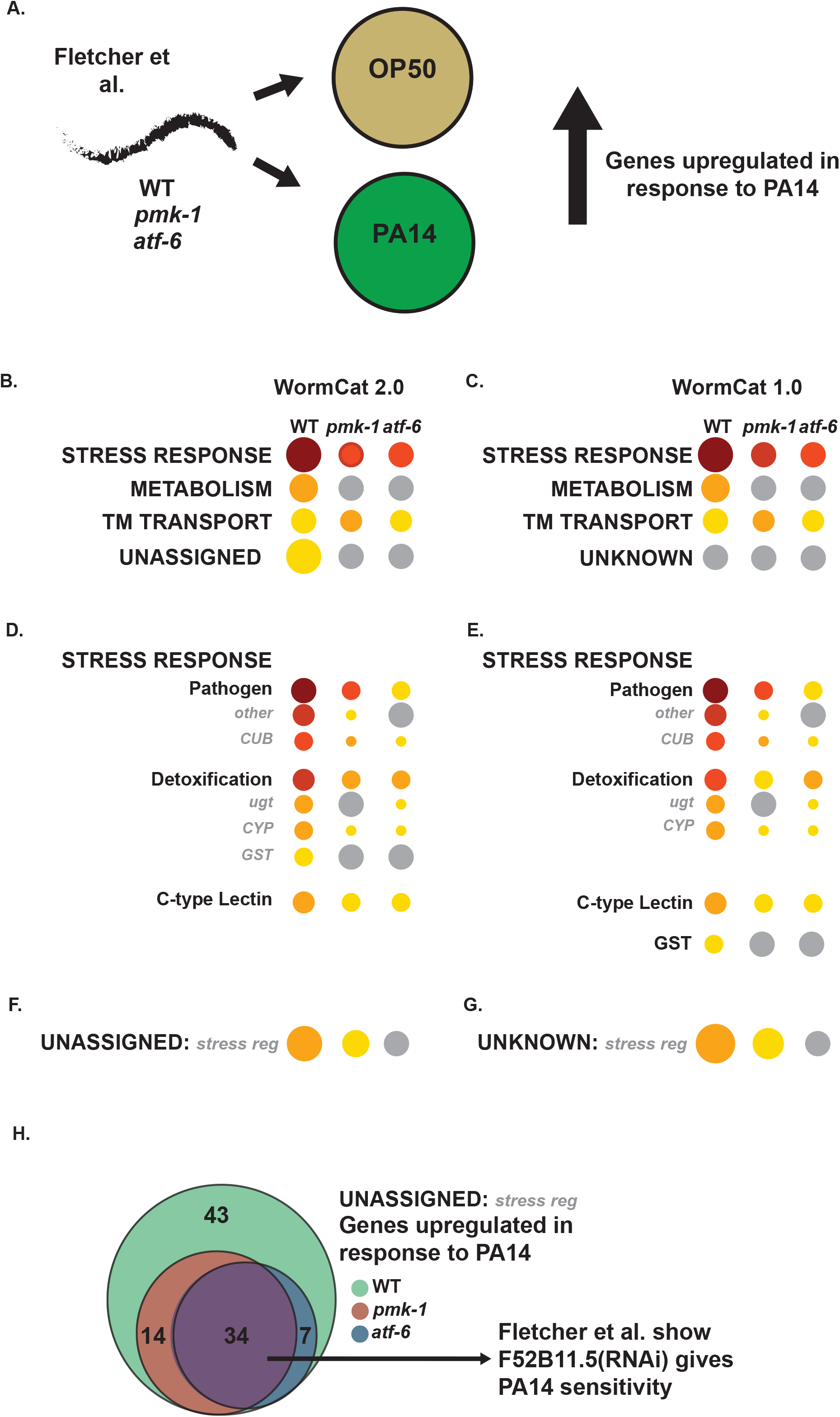
UNASSIGNED: regulated by multiple stress genes are regulated by a *pmk-1/atf-7* immunity circuit in response to *Pseudomonas aeruginosa* (PA14). **A**. Schematic diagram showing experimental workflow from Fletcher et al. investigating gene regulation in response to PA14. Cat1 category enrichment in WormCat 2.0 (**B)** compared with WormCat 1.0 (**C**). Cat2 level enrichment of Stress Response and Unassigned/Unknown in WormCat 2.0 (**D, F**) and WormCat 1.0 (**E, G**). **H**. Venn Diagram showing dependence on *pmk-1* and *atf-7* of UNASSIGNED genes upregulated by PA14. See also **Table S5**. Abbreviations: TM, transmembrane; CYP, cytochrome p450; CUB, complement C1r/C1s, Uegf, Bmp1; ugt, uridine diphosphate glucuronosyl transferase; GST, Glutathione S-transferase.

Interestingly, many of the MSR Genes upregulated in response to PA14 are also dependent on *atf-7* and the *pmk-1* pathway (Figure 4F-H; Supplemental Table 5). Fletcher et al. tested 45 PA14 upregulated/*atf-7* dependent genes for PA14 sensitivity and found 14 survived less well on pathogenic bacteria (Esp phenotype) (Fletcher et al., 2019). Many of these genes with the Esp phenotype contained domains known to be important for pathogen responses. However, the authors found an Esp phenotype for one ATF-7-target with no known domains (Fletcher et al., 2019). This gene, F52B11.5, is a WormCat MSR gene and is also regulated by methylmercuric chloride, Cry5B, and *hif-1/HIF1*. Thus, genes within the MSR gene category appear to be regulated similarly to genes in well-described pathogen responses, including genes with critical biological responses. By defining a category for enrichment, WormCat provides a framework for future studies of genes that would ordinarily be overlooked for further analysis because less is known about their function.

### Tissue-specific expression of UNASSIGNED genes

Identification of category enrichment commonly depends on Fisher’s exact test, a statistical metric that builds a contingency table to determine the likelihood that the number of items in the group in a test set is more enriched than the number of those items in the entire set. Thus, enrichment statistics can be affected by the number of items in the entire set. We sought to explicitly test the hypothesis that excluding genes based on annotation status altered the statistical metrics using a tissue-specific RNA-seq data set published by the Ahringer lab (Serizay, et al). We determined category enrichment RNA-seq data from two tissues, Intestine-only and Neurons-only, and compared two versions of the WormCat annotation list, All (including UNASSIGNED genes) and Assigned only. While the top categories of STRESS RESPONSE, PROTEOLYSIS PROTEOSONE, TRANSCRIPTION FACTOR, and TRANSMEMBRANE TRANSPORT remained significant, METABOLISM failed significance at the FDR correction (Figure 5B, Supplemental Figure 3A, B; Supplemental Table 6) although the *C. elegans* intestine has a clearly defined role in metabolism (Dimov and Maduro, 2019; McGhee, 2007). At the Category 2 level, removing the UNASSIGNED genes alters enrichment of detoxification genes in the STRESS RESPONSE category for intestinally expressed genes; however, METABOLISM: Lipid remains statistically enriched (Figure 5B, Supplemental Figure 3A, B; Supplemental Table 6).

**Figure 5:**
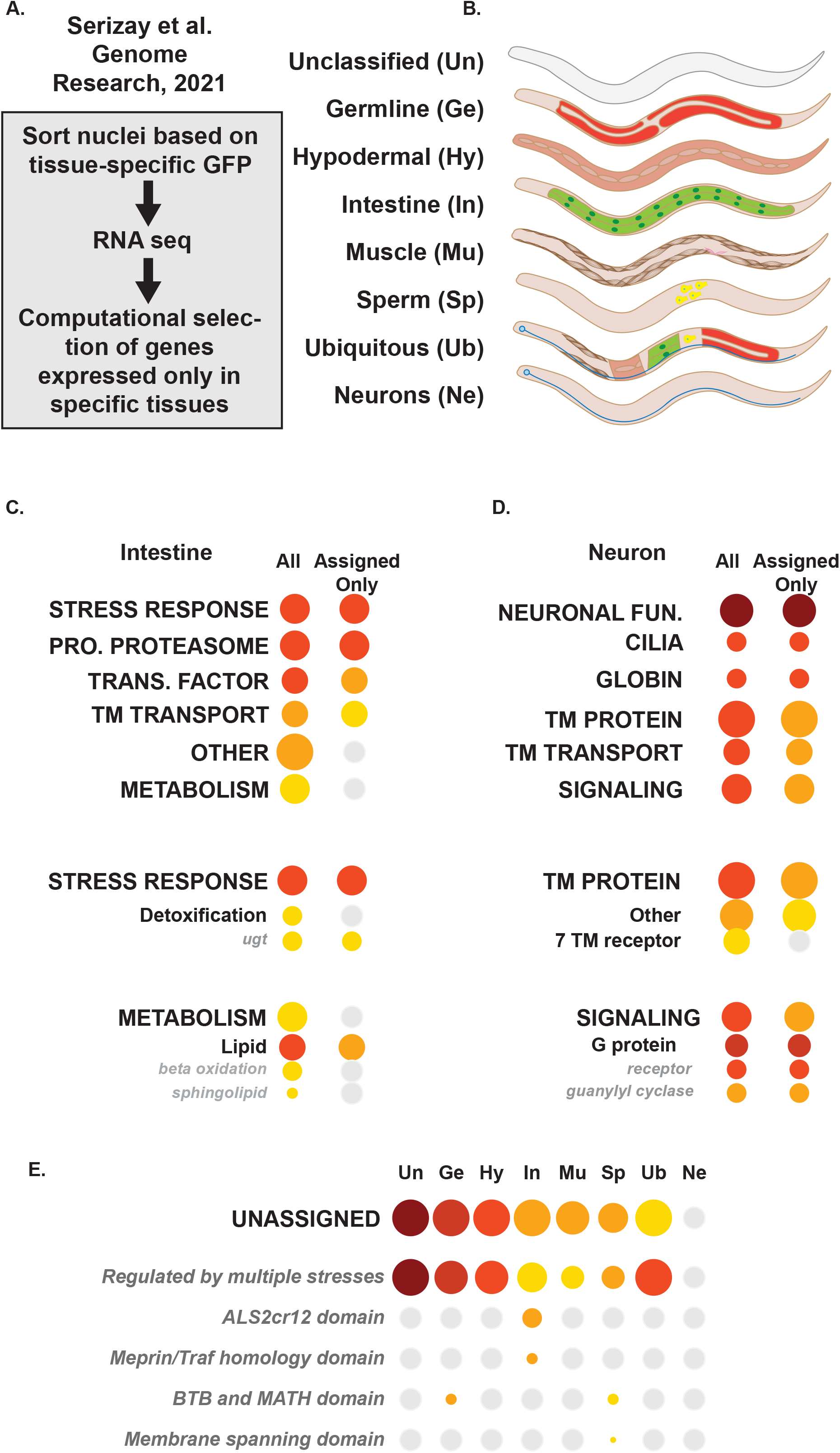
Tissue-specific RNA-seq shows enrichment of UNASSIGNED genes in data sets specific to multiple cell types, but not neurons. **A**. Schematic diagram showing workflow from Serizay et al. Comparison of category enrichment with inclusion (All) and exclusion (Assigned only) of UNASSIGNED genes in Intestine-only (**B**) and Neuron-only (C) genes. **E**. Cat3 breakdown of UNASSIGNED genes in Unclassified (Un), Germline (Ge), Hypoderm (Hy), Intestine (In), Muscle (Mu) Sperm (Sp), and Neuron-only genes. See also **Supplemental Table 4** and **Table S6**. Abbreviations: Pro, Proteolysis; Trans., Transcription; TM, transmembrane; ugt, uridine diphosphate glucuronosyl transferase.

On the other hand, UNASSIGNED genes are not enriched in Neuronal-only RNA-seq data sets, and the major enriched Cat1 groups all remain statistically significant (Figure 5B, C; Supplemental Figure 3C; Supplemental Table 6). At the Cat2 level, removing the UNASSIGNED genes shifts the 7TM protein out of the significance range, but SIGNALING Cat2 groups remain significant. Taken together, this evaluation shows that excluding unannotated genes has different effects on pathway analysis of RNA-seq from distinct tissues. Thus the UNASSIGNED gene category has an important role in stabilizing effects on annotation bias.

The observation that there were different numbers of UNASSIGNED genes in Intestine-only vs. Neuronal-only tissues in the Serizay, et al. data prompted us to examine enrichment of UNASSIGNED genes in other tissues in this data set. Strikingly, Neuronal-only tissue is the only group that lacks enrichment of UNASSIGNED genes (Figure 5D; Supplemental Table 6). This appears to be largely driven by the UNASSIGNED MSR genes. One explanation for the difference in tissue distribution of the UNASSIGNED genes could be that genes expressed in neurons are more likely to be conserved, well-studied, and better annotated. In order to address this question, we examined the numbers of GO annotated genes and predicted human orthologs in our NEURONAL FUNCTION category. These genes were defined as having curated functions in neurons that were not shared with other tissues, and NEURONAL FUNCTION is enriched in multiple independent tissue-specific datasets (Holdorf, et al. 2020; Figure 5D). WormCat NEURONAL FUNCTION genes were well-annotated by GO, and about half the genes had predicted human orthologs in the Parasite Biomart (Figure 2D, E; Table S2A, Supplemental Table 6). However, nuclei isolated from neurons by Serizay et al show enrichment in categories with low numbers of human orthologs, such as TM transport and TM protein (Figure 2I; Figure 5D); therefore, there may be other explanations for the apparent tissue-specificity of the UNASSIGNED genes.

### UNASSIGNED genes are poorly enriched in proteomics samples

The annotation lists in WormCat were designed for whole-genome RNA-seq or ChIP-seq assays and so included both protein-coding and non-coding genes (Holdorf, et al. 2020). To provide an appropriate annotation list for proteomics, we removed non-transcribed genes (ORF only annotation list). This annotation list retains the category of NON-CODING RNA; however, the genes that remain function in processing non-coding RNAs (Table S1). Next, we examined gene enrichment patterns in two published proteomics datasets (Narayan et al., 2016; Reinke et al., 2017). Reinke et al. performed proteomics on nuclear vs. cytoplasmic fractions as well as obtaining nuclear and cytoplasmic proteomes from multiple tissues (Reinke et al., 2017) see Figure 6A).

**Figure 6:**
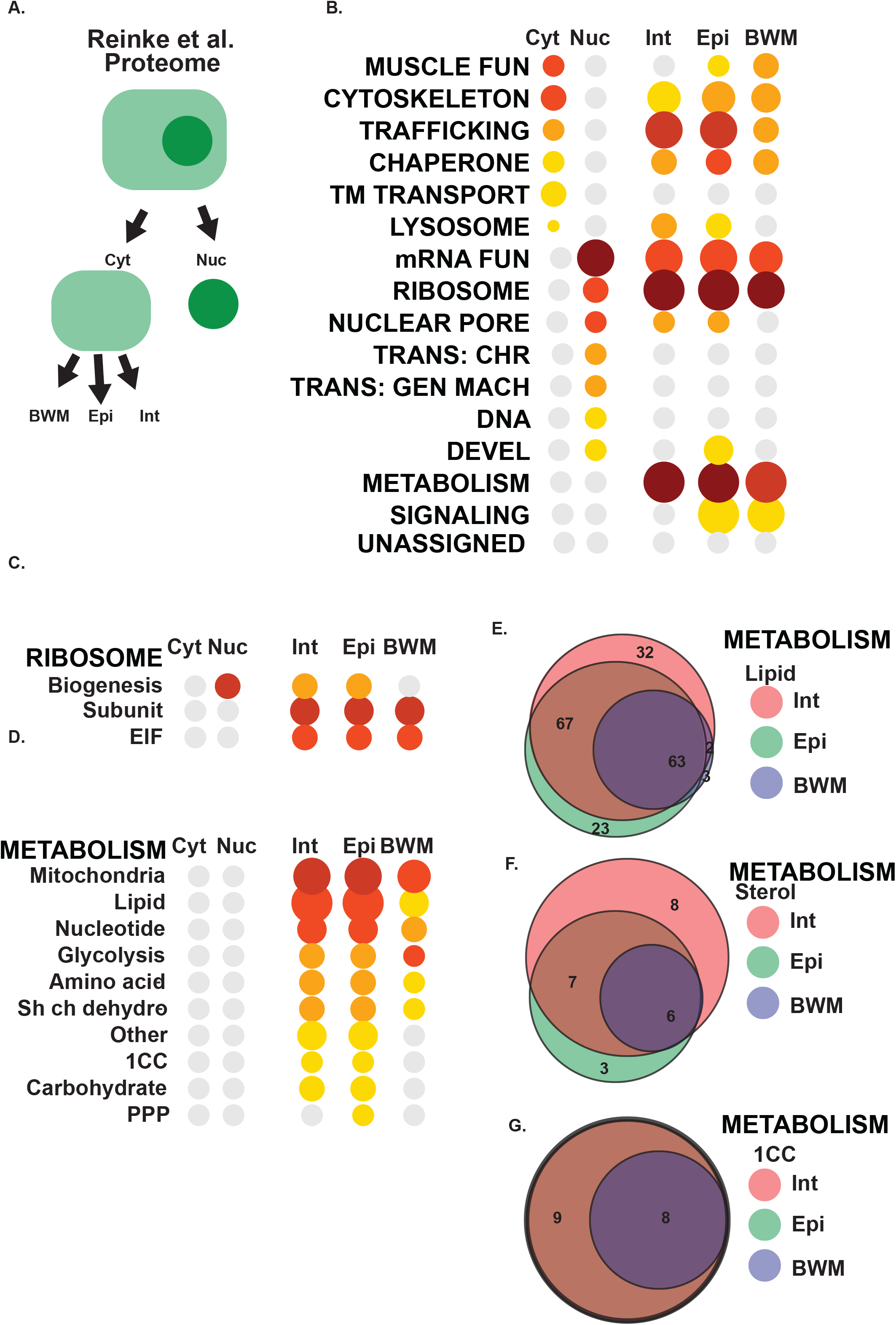
Tissue and compartment-specific proteomic data lack enrichment in UNASSIGNED genes. **A**. Workflow for Reinke et al. **B**. Cat1 level enrichment of proteins in cytoplasmic or nuclear extracts along with cytoplasmic extracts from Intestine (Int), Epidermis (Epi), or Body Wall Muscle (BWM). Breakdown of WormCat enrichment in Ribosome (**C**) or Metabolism (**D**). Venn Diagrams showing the number of tissue-specific genes in METABOLISM (**E**), METABOLISM: Lipid (**D**), or METABOLISM: 1CC). See also **Supplemental Figure 5** and **Table S7**. Abbreviations: Cyt, Cytoplasmic; Nuc; Nuclear, Trans., Transcription; TM, transmembrane; EIF, elongation and initiation factor; 1CC, 1-carbon cycle; PPP, pentose phosphate pathway.

Enrichment of mRNA functions, NUCLEAR PORE, and the Transcription associated categories in the nuclear fractions were expected; however, RIBOSOME was strongly enriched (Figure 6B; Supplemental Table 7). As ribosomes are localized to the cytoplasm, we examined the gene enrichment patterns at the Cat2 level and found the major enrichment in the nuclear samples was in RIBOSOME: biogenesis, which is a nucleolar process (Figure 6C; Supplemental Table 7). RIBOSOME: Subunit and RIBOSOME: EIF were strongly enriched in the cytoplasmic fractions of intestinal, epidermal, and body wall muscle cells. Comparison of category enrichment between cell types did not reveal large differences at the Cat1 level (Figure 6B; Supplemental Table 7). Both the intestine and epidermis play important roles in in *C. elegans* metabolism. Detailed examination of these proteomes also showed similar patterns (Figure 6C; Supplemental Table 7). We next asked if the same proteins were driving enrichment in the METABOLISM subcategories or if there were tissue-specific differences. While many of the proteins in the category “METABOLISM: Lipid” were shared, some were tissue-specific (Figure 6E; Supplemental Table 7). The Cat3 group with the most differences was METABOLISM: Lipid: sterol (Figure 6F; Supplemental Table 7). The group includes many short-chain dehydrogenases that can contribute to hormone production (Mahanti et al., 2014) and thus have may tissue-specific functions. In contrast, the 1 Carbon-cycle (1CC) genes detected in the proteomics were the same in the intestine and epidermis (Figure 6G; Supplemental Table 7).

In contrast to multiple tissue-specific RNA-seq data sets (Figure 5; see also Holdorf et al. 2020), the UNASSIGNED category is not enriched in any of the samples in the Reinke et al. data. In order to verify this in a different proteomics data set, we used the ORF annotation list in WormCat to find enriched categories in proteomics comparing young adult and aging worms (Narayan et al., 2016). The authors also performed fractionations separating cytoplasmic, membrane-bound, and nuclear proteins (Figure S4C; Supplemental Table 8). Because of the lower number of detected membrane-bound and nuclear proteins, we limited our search for enriched categories to “all detected”, “changed in aging” and the “cytoplasmic fraction”. WormCat finds strong enrichment of METABOLISM, mRNA FUNCTIONS, and RIBOSOME in the cytoplasmic category, whereas enrichment DNA functions and NUCLEAR PORE are lost in comparison to All Detected (Figure S4D; Supplemental Table 8). METABOLISM: Lipid and STRESS RESPONSE are enriched in proteins that change during aging (Figure S4D; Supplemental Table 8). Similar to the proteomics data from Reinke, et al., there is no enrichment for the UNASSIGNED category. As the authors of Reinke et al. point out, proteomics is less sensitive than RNA seq, and several studies have pointed out a discordance between mRNA and protein expression (Harvald et al., 2017). Thus, proteins in the UNASSIGNED group could be lower than the threshold for proteomics detection.

## Discussion

### Strategies for study of poorly annotated genes

Fully sequenced genomes for the major model systems as well as humans have been available for 20 or more years (Gates et al., 2021). However, each of these genomes contains large numbers of genes that are poorly annotated. For example, the human genome contains around 20,000 protein-coding genes, and 3,000 have limited functional annotations (Dey et al., 2015). Even well-characterized eukaryotes with smaller genomes, such as *Saccharomyces cerevisiae* contain large numbers of genes with unknown functions (Wood et al., 2019). These genes may be understudied due to research bias if no obvious human ortholog or disease relevance is evident. The biological functions of these genes may be elusive because of functional redundancy or because they function in specific contexts that are difficult to replicate in experimental environments. One approach to providing functional information on these genes is to determine mRNA expression patterns and define co-expression groups (Pertea et al., 2018) to identify processes that might be shared among the co-expressed genes. Other studies have used protein: protein interactions defined by either yeast two-hybrid (Y2H) (Rolland et al., 2014) or mass spec (Huttlin et al., 2017) to link poorly annotated proteins to well-described processes. Indeed, Roland and Vidal found that the accumulation of Y2H data increased study of and publications on individual genes (Rolland et al., 2014). Thus, providing interaction or co-expression data on poorly annotated genes can spur studies that provide insight into biological function.

The *C. elegans* genome contains around 19,000 protein-coding genes (Schwarz, 2005). Large-scale RNAi and mutant screens ascribe phenotypes to around 1/3 of the genome (Schwarz, 2005), and other genes may be annotated based on homology to human genes (WormBase). However, 26% of the genomes lack GO annotation (Figure 2A). Early estimates projected that 9% of the *C. elegans* genome is conserved across metazoa, yet the functions of these genes are unknown (Schwarz, 2005). The exclusion of poorly annotated genes not only biases statistical metrics used in enrichment analysis (see Figure 4, Figure S4, Table S5) but also discourages further study of these genes. By improving annotation of UNKNOWN/UNASSIGNED genes, the analysis provided by WormCat 2.0 allows these genes to appear in enrichment analysis. Using an RNA-seq data set exploring the bacterial pathogen response (Fletcher et al., 2019), we found that more than half of the enriched UNASSIGNED: multiple stress-regulated genes were regulated by the same signaling and transcriptional network that controlled the canonical stress response (Figure 3). Interestingly, the authors found that one of these genes affected survival upon PA14 exposure (Fletcher et al., 2019), demonstrating biologically relevant activities in this gene set.

The genes in the UNASSIGNED set have a higher proportion that lack GO terms, and many appear to lack human orthologs or are lineage-specific. Some of these genes may have homology that is missed due to structural similarity that is not reflected at the amino acid level or because homology is undetectable by BLAST (Pearson, 2013). We used abSENSE, a recently developed tool that uses evolutionary distances to determine whether a gene could appear lineage-specific merely because of failure to detect homologs in outgroups. We found that homology detection failure could be sufficient to explain lack of orthologs in other *Caenorhabditis* species, the clade III nematodes *Brugia malayi* and *Loa loa* and a non-nematode invertebrate, the echinoderm *Strongylocentrotus purpuratus*. We found that most UNASSIGNED genes were found to have orthologous proteins in *S. purpuratus* and so were not lineage-specific. For others, however, homology detection failure appeared to be a plausible explanation for the apparent lineage-specificity. This result indicates that the number of lineage-specific genes may be overestimated (Weisman et al., 2020). UNASSIGNED genes that appear *C. elegans*-specific may not be selected for functional testing as they lack definable human disease or functional relevance. It is striking that the UNASSIGNED genes are underrepresented in neuronal cells in published tissue-specific RNA-seq data from Serizay, et al. (See figure). This could be an artifact of our annotation strategy; however, it might also reflect tighter evolutionary constraints in these specialized cells.

### Strategies for category enrichment

The tractability and affordability of -omics technologies have allowed *C. elegans* researchers to compare whole-genome mRNA or protein distributions between mutant or RNAi backgrounds and a wide variety of environmental conditions. These studies then rely on category enrichment tools to identify genes for further analysis. We have extensively compared WormCat to GO-based enrichment tools (Holdorf, et al, 2020) and found that WormCat identifies biologically relevant gene sets not revealed in GO. However, alternative category enrichment tools employ distinct strategies, and dataset analysis may benefit from cross-platform analysis. For example, EVITTA, a web-based tool for RNA seq analysis, allows downloads from GEO and enrichment by KEGG, GO, or WormCat term along with several alternatives for visualization (Cheng et al., 2021). The Kenyon lab developed the Gene Modules tool, which integrates gene co-expression data (Cary et al., 2020). Functional characterization of poorly annotated genes is a complex problem that may require multiple strategies to reveal gene-phenotype connections. *C. elegans* is an excellent model for determining such gene phenotype interactions. WormCat 2.0 provides a platform that allows characterization of understudied genes, thus opening up a previously enigmatic group of *C*.*elegans* genes for further study.

## Methods

### Annotations

WormBase version WS270 was utilized for WormBase descriptions (https://wormbase.org). We used ParaSite Biomart to obtain GO terms and predicted human orthologs for *C. elegans* genes (https://parasite.wormbase.org/biomart/martview). NCBI Conserved Domain Database (https://www.ncbi.nlm.nih.gov/Structure/cdd/wrpsb.cgi) and TOPCONS (https://topcons.cbr.su.se) were used for membrane-spanning domain predictions.

### abSENSE

abSENSE was downloaded from the publicly available github, https://github.com/caraweisman/abSENSE, and run locally. Run parameters were (info from run file). The input bitscore file was (derived from/can be found in/etc). The input evolutionary distance file was derived from 10 eukaryotic BUSCO genes determined to be single-copy in all analysis species, which were then aligned using MUSCLE (default parameters) and used as input to the PROTDIST program (default settings) from the PHYLIP package.

### Scripts

WormCat 2.0 is built on the foundations of the original WormCat Web python codebase. A key new feature of Wormcat 2.0 is “Batch Processing,” which allows the execution of multiple datasets using a single submission. To support batch processing, we leveraged several open-source python packages and tools (Pandas, Celery & Redis). The Pandas package handles data and file processing while Celery is used to queue and distribute the batch tasks, and Redis is the in-memory data structure store used as a message broker for Celery. All three of these technologies use the Open Source BSD license.

### Data Availability

The WormCat 2.0 code is available under MIT Open Source License and can be downloaded from the GitHub repository https://github.com/dphiggs01/wormcat along with all annotation lists and version-control information. WormCat can also be installed R package using the devtools library for direct usage.

## Acknowledgments

We wish to thank members of the Walker lab, Dr. Micheal Lee and Dr. Marian Walhout (UMASS Chan School of Medicine) for their helpful discussion. We are also grateful to Life Science Editors and Dr. Sabbi Lall for manuscript editing. Funding to AKW NIH NIA 1R01AG053355.

## Figure Legends

**Figure S1: Unassigned genes in *C. elegans* include a subset with human orthologs. A**. Breakdown of WormCat Cat1 level categories with numbers of genes annotated by GO. Cat2 GO breakdown of TM protein/Signaling (**B**) and Stress Response at the Cat2 (**C**) and Cat3 level (**D**). **E**. Breakdown of WormCat Cat1 level categories with numbers of genes designated as having human orthologs in the ParaSite Biomart. Cat2 human ortholog breakdown of TM protein/Signaling (**F**) and Stress Response at the Cat2 (**G**) and Cat3 level (**H**). Red percentages denote categories with substantially more human orthologs in a genome-wide BLASTP comparison of each *C. elegans* gene with the human genome (see **Table S4**). Abbreviations: TM, transmembrane; Stress resp, Stress response; Prot prot, Proteolysis Proteasome; Trans factor, Transcription factor; EC, Extracellular; mRNA fun, mRNA function, Prot general, Proteolysis General; Neuro fun, Neuronal function; Protein mod, Protein modification; Devel, Development; Trans chr, Transcription Chromatin; Trans. GM, Transcription: General Machinery; Sig. GPCR, Signaling: G-protein coupled receptor; CUB, CUB, complement C1r/C1s; NLP, neuropeptide-like protein; CYP, cytochrome p450; ugt, uridine diphosphate glucuronosyl transferase; GST, Glutathione S-transferase.

**Figure S2: Unassigned genes in *C. elegans* include predicted lineage-specific and non-lineage-specific genes. A**. Pie chart of *C. elegans* protein-coding genes predicted to be lineage-specific by Zhou et al (Zhou et al., 2015). **B**. Breakdown of WormCat Cat1 level categories with numbers of predicted lineage-specific genes. **C**. Cat2 GO breakdown of Unassigned genes. **D**. Venn Diagram illustrating the number of lineage-specific genes and those with no predicted human ortholog within the UNASSIGNED gene set. Predicted lineage-specific gene number Cat2 level categories in Neuronal Function (**E**) and Cat3 level categories in Synaptic Function (**F**). Breakdown of numbers of lineage-specific genes within the Cat2 level of TM protein/Signaling (**G**) and Stress Response at the Cat2 (**H**) and Cat3 level (**I**). (See **Table S4**). Abbreviations: TM, transmembrane; msr, multiple stress-regulated; mem span, membrane-spanning; GT family A, glucosyltransferase family A; TTR, TransThyretin-Related family domain; BTB/MATH, BR-C, ttk, and bab/meprin and TRAF homology; Synaptic fun, Synaptic function; Devel, Development; Trans, Transcription; NP, Neuropeptide, NT receptor, Neurotransmitter receptor; NT met, Neurotransmitter metabolism; Vesicle traff, Vesicle traffic; Stress resp, Stress response; Prot prot, Proteolysis Proteasome; Trans factor, Transcription factor; EC, Extracellular; mRNA fun, mRNA function, Prot general, Proteolysis General; Neuro fun, Neuronal function; Protein mod, Protein modification; Devel, Development; Trans chr, Transcription Chromatin; Trans. GM, Transcription: General Machinery; Sig. GPCR, Signaling: G-protein coupled receptor; CUB, CUB, complement C1r/C1s; NLP, neuropeptide-like protein; CYP, cytochrome p450; ugt, uridine diphosphate glucuronosyl transferase; GST, Glutathione S-transferase.

**Figure S3: Contingency tables comparing the significance of Cat1 level categories with and without UNASSIGNED genes**. Data tables showing numbers of genes and enrichment values when all genes (**A, C**) are used rather than functionally annotated genes only (Assigned only) (**B, D**) in the intestine (**A, B**) and Neuron (**C, D**). the p-value is Fisher’s Exact Test; gray highlighting shows values that are below an FDR of 0.01. Abbreviations: TM, transmembrane; RGS, Regulated Gene Set.

**Figure S4 Tissue and compartment-specific proteomic data lack enrichment in UNASSIGNED genes. A**. Workflow for Reinke et al. **B**. Cat1 level enrichment of proteins in cytoplasmic or nuclear extracts along with nuclear extracts from Intestine (Int), Epidermis (Epi), or Body Wall Muscle (BWM). **C**. Workflow schematic for Narayan, et al. D. Cat1 level enrichment of proteins from all conditions, those that are changing in aging animals (Δ Aging) and from Cytoplasmic extracts. See also **Table S7, Table S8**. Abbreviations: TRANS., transcription; PROT., proteolysis; SILAC, Stable Isotope Labeling by/with Amino acids.

**Supplemental Table 1: WormCat annotations**. xlsx file containing annotation definitions.

**Supplemental Table 2: WormCat annotation definitions**. xlsx file containing annotation definitions. Tabs: 1. Cat1 definitions, 2. Cat2 definitions, 3. Cat3 definitions. Color key: light blue highlight, other to assigned; light yellow highlight, new category; border, substantial change of genes in a category.

**Supplemental Table 3: WormCat ORF annotations**. xlsx file containing annotation definitions for protein-coding genes.

**Supplemental Table 4: GO terms and human orthologs for WormCat genes**. xlsx file matching WormCat protein-coding genes to GO terms and human orthologs. Tabs: 1. GO_human_ortholog from, 2. human.bit_abSENSE, 3. Unassigned_GoRILLA, 4. GoRILLa_listed_gene.GO.

**Supplemental Table 5: WormCat output for Fletcher at al**.. xlsx file comparing WormCat 2.0 and WormCat 1.0 annotation lists. Blue shading show significance level. Tabs: 1. WC2.0_Cat1, 2. WC2.0_Cat2, 3. WC2.0_Cat3, 4. N2_up_WC2.0_cat, 5. up_pmk-1_WC2.0_cat6. up_atf-7_WC2.0_cat, 7. N2_down_WC2.0_cat, 8. down_pmk-1_WC2.0_cat, 9. down_atf-7_WC2.0_cat, 10. WC1.0_Cat1, 11. WC1.0_Cat2, 12. WC1.0_Cat3, 13. N2_up_WC1.0_cat, 14. up_atf-7_WC1.0_cat, 15. up_pmk-1_WC1.0_cat, 16. N2_down_WC1.0_cat, 17. down_pmk-1_WC1.0_cat, 18. down_atf-7_WC1.0_cat.

**Supplemental Table 6: WormCat output for Serizayat al**.. xlsx file comparing WormCat 2.0 annotation lists with and without UNASSIGNED genes. Blue shading shows the significance level. Tabs1. WC2.0_all_Cat1, 2. WC2.0_all_Cat2, 3. WC2.0_all_Cat3, 4. WC2.0_Assingned_Cat1, 5. WC2.0_Assingned_Cat2, 6. WC2.0_Assingned_Cat3, 7. Contingency_All, 8. Contingency_intestine, 9. Contingency_neurons.

**Supplemental Table 7: Protein category enrichment in Reinke et al**. xlsx file containing WormCat output for proteomics using the ORF only annotation list. Blue shading shows the significance level. Tabs: 1. Cat1, 2. Cat2, 3. Cat3, 4. Cytoplasm-specific genes,5. Nucleus-specific genes,6. Body wall muscle (BWM) cytoplasmic genes,7. Epidermis cytoplasmic genes,8. Intestine cytoplasmic genes,9. Body wall muscle (BWM) nuclear genes,10. Epidermis nuclear genes, 11. Intestine nuclear genes.

**Supplemental Table 8: Protein category enrichment in Narayan et al**. xlsx file containing WormCat output for proteomics using the ORF only annotation list. Blue shading shows the significance level. Tabs: 1. Cat1, 2. Cat2, 3. Cat3, 4. all_detected_peptides_cat, 5. aging.change_cat, 6. Cytoplasm_cat.

