## Supplemental Figures 1-4 for "WormCat 2.0 defines characteristics and conservation of poorly annotated genes in *Caenorhabditis elegans*"

Higgins, et al.
Supplemental Figure 1

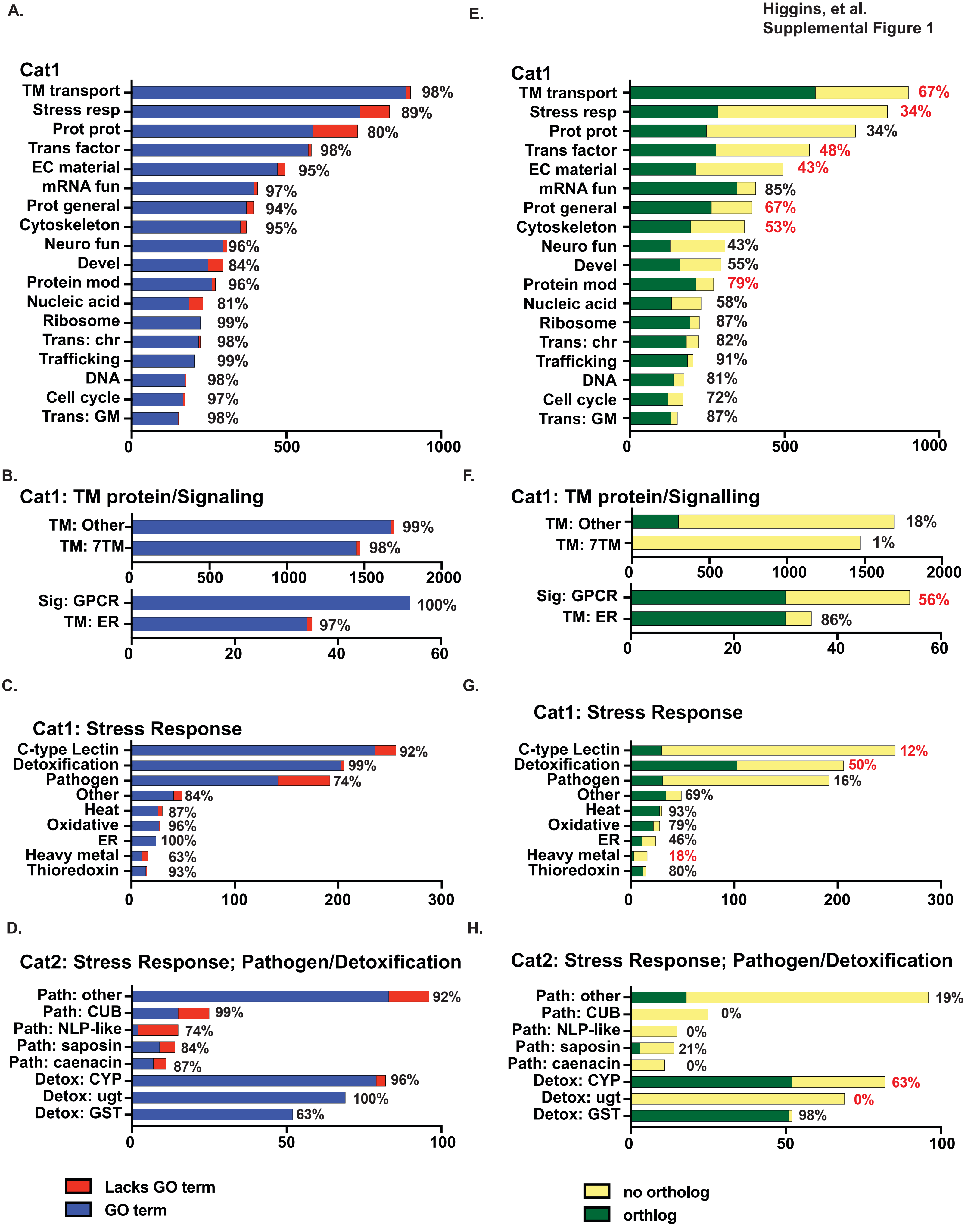

Higgins, et al.
Supplemental Figure 2

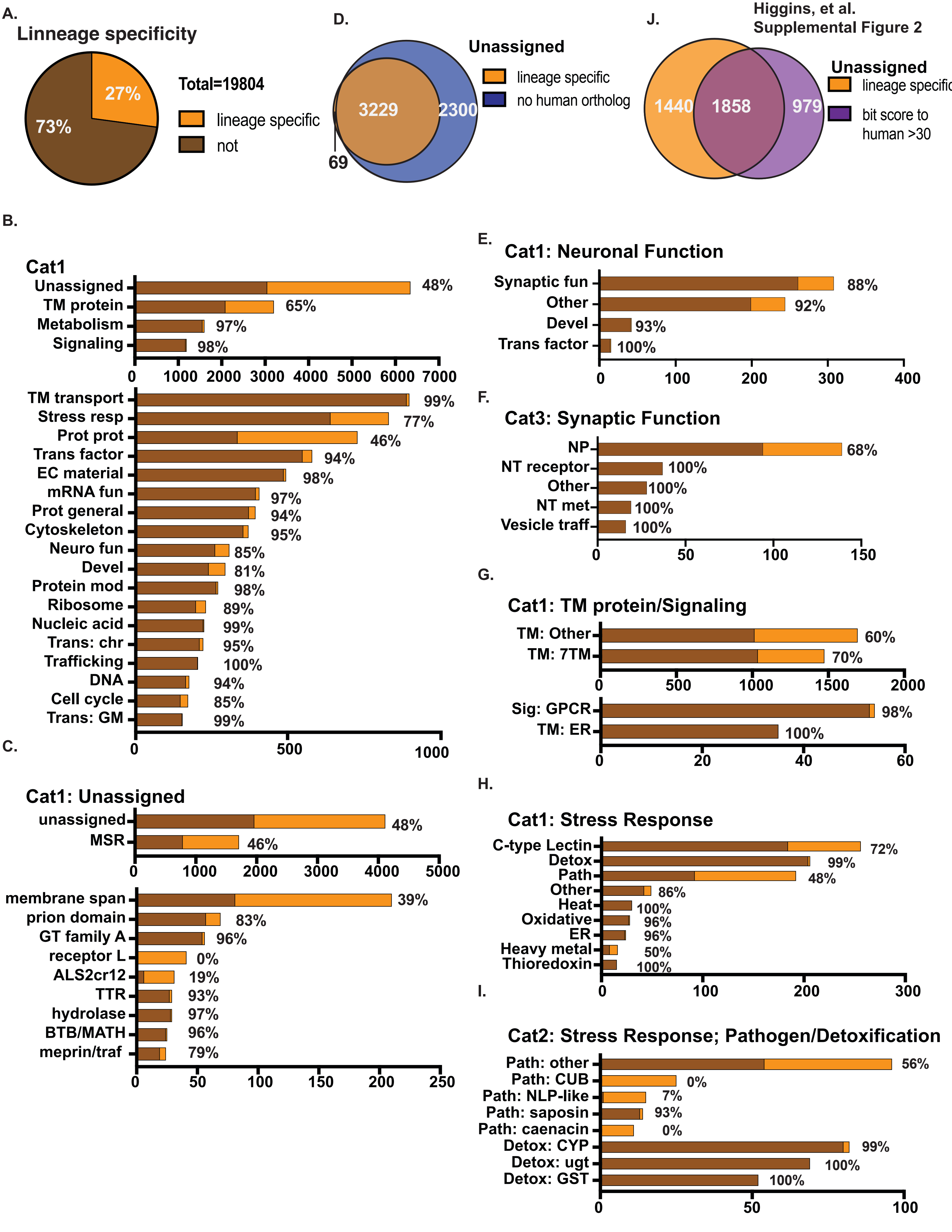

**A.**

| Category 1 | All | Intestine_only_RGS | *p value* |
|---|---|---|---|
| Stress response | 833 | 91 | 1.91E-21 |
| Proteolysis proteasome | 730 | 81 | 1.01E-19 |
| Transcription factor | 581 | 64 | 6.51E-16 |
| TM transport | 901 | 69 | 2.63E-10 |
| Metabolism | 1601 | 81 | 3.59E-10 |
| Unassigned | 6343 | 303 | 6.30E-05 |
| Total genes | 31389 | 964 | |

**B.**

| Category 1 | Assigned only` | Intestine_only_RGS | *p value* |
|---|---|---|---|
| Stress response | 833 | 91 | 1.06E-15 |
| Proteolysis proteasome | 730 | 81 | 1.50E-14 |
| Transcription factor | 581 | 64 | 8.26E-12 |
| TM transport | 901 | 69 | 8.90E-07 |
| Metabolism | 1601 | 81 | 0.017721 |
| Total genes | 25046 | 661 | |

**C.**

| Category 1 | All | Neurons_only_RGS | *p value* |
|---|---|---|---|
| Neuronal function | 308 | 149 | 2.28E-111 |
| Cilia | 60 | 25 | 1.85E-19 |
| Globin | 36 | 19 | 1.21E-16 |
| TM protein | 3200 | 176 | 3.13E-15 |
| TM transport | 901 | 74 | 5.79E-15 |
| Signaling | 1188 | 84 | 2.14E-13 |
| Unassigned | 6343 | 198 | 0.035747 |
| Total genes | 31389 | 833 | |

**D.**

| Category 1 | Assigned only` | Neurons_only_RGS | *p value* |
|---|---|---|---|
| Neuronal function | 308 | 149 | 3.05E-98 |
| Cilia | 60 | 25 | 3.00E-17 |
| Globin | 36 | 19 | 6.18E-15 |
| TM protein | 3200 | 176 | 1.35E-10 |
| TM transport | 901 | 74 | 6.21E-09 |
| Signaling | 1188 | 84 | 2.48E-08 |
| Total genes | 25046 | 635 | |

Higgins, et al.
Figure S4

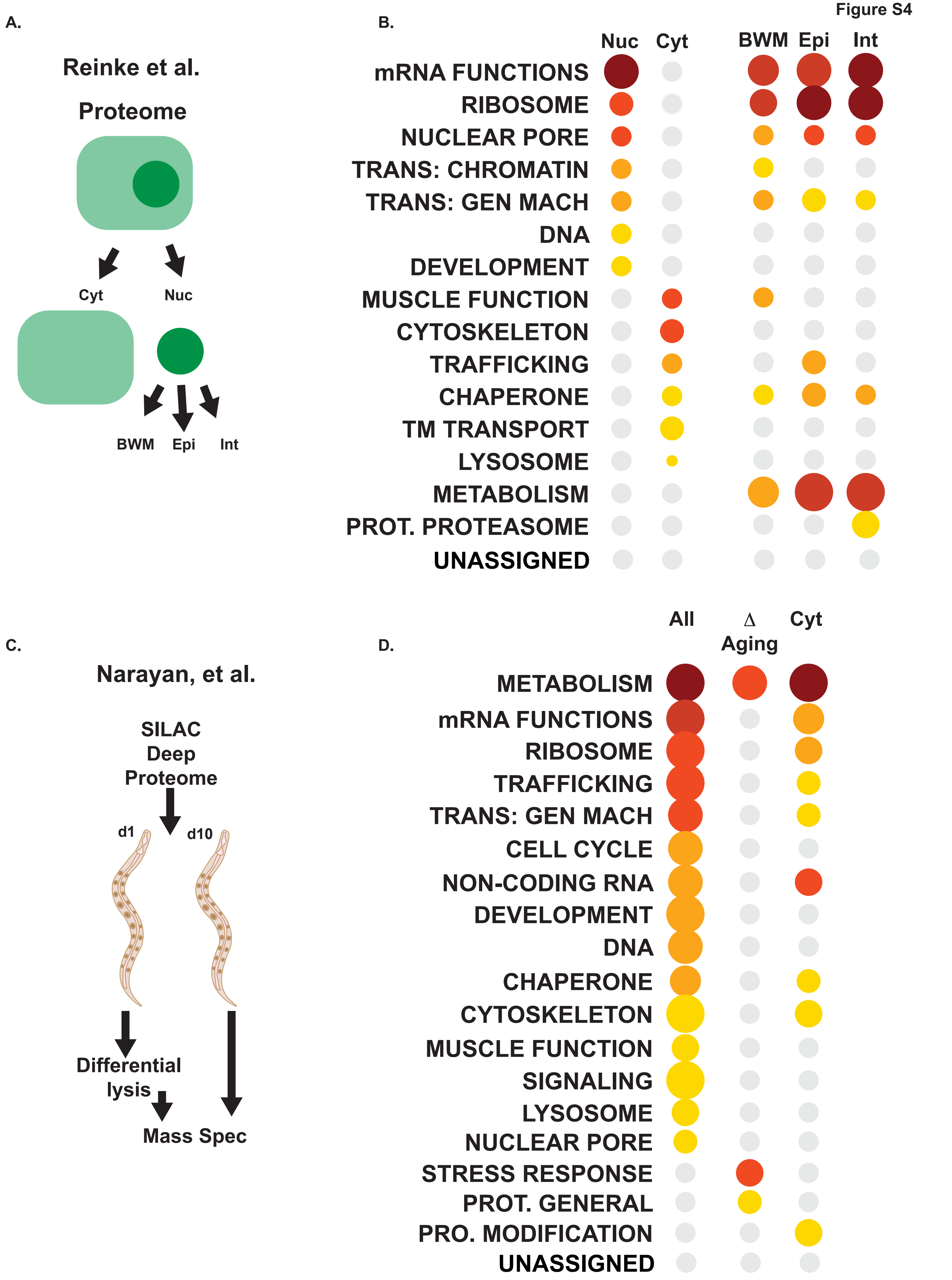
